# Novel clock neuron subtypes regulate temporal aspects of sleep

**DOI:** 10.1101/2024.05.03.592478

**Authors:** Dingbang Ma, Jasmine Quynh Le, Xihuimin Dai, Madelen M. Díaz, Katharine C. Abruzzi, Michael Rosbash

## Abstract

Circadian neurons within animal brains orchestrate myriad physiological processes and behaviors, But the contribution of these neurons to the regulation of sleep is not well understood. To address this deficiency, we leveraged single-cell RNA sequencing to generate a new and now comprehensive census of transcriptomic cell types of *Drosophila* clock neurons. We focused principally on the enigmatic DN3s, which constitute about half of the 75 pairs of clock neurons in the fly brain and were previously almost completely uncharacterized. These DN3s are organized into 12 clusters with unusual gene expression features compared to the more well-studied clock neurons. We further show that different DN3 subtypes with distinct projection patterns promote sleep at specific times of the day through a common G protein–coupled receptor, *TrissinR*. Our findings indicate an intricate regulation of sleep behavior by clock neurons and highlight their remarkable diversity in gene expression, projection patterns and functional properties.

## Introduction

A comprehensive characterization of transcriptomic neuron types, their projection patterns and how they give rise to different complex behaviors are major goals of contemporary neuroscience research. The circadian clock and its brain neurons are ideal substrates to help achieve these aims (*1, 2*). The clock is regulated by a transcription-translation feedback loop, which resides within these brain clock neurons and times different rhythmic behaviors of animals (*3, 4*).

The principal circadian pacemaker of mammals is localized in the suprachiasmatic nucleus (SCN) in the brain. Each unilateral SCN of the mice brain contains about 10,000 neurons, which has been classically divided into two main regions based on the expression of two neuropeptides: the vasoactive intestinal polypeptide (VIP) expressing core and arginine vasopressin (AVP) expressing shell (*5*). Recent single cell RNA sequencing studies have revealed even more heterogeneity. There are for example at least five different neuronal cell types in the SCN (*6–8*). Relevant to sleep regulation is the innervation of the dorsomedial hypothalamic nucleus (DMH) by SCN processes: The DMH inhibits the activity of sleep-promoting ventrolateral preoptic (VLPO) neurons and activates the activity of wake-promoting hypocretin neurons via different neurotransmitters (*9–12*).

In *Drosophila*, there are approximately 150 clock neurons in total. Based on the anatomical locations of these cell bodies and their projection patterns, there are at least four dorsal neurons groups (DN1a, DN1p, DN2, DN3) as well as five lateral neuron groups. The latter includes small ventral lateral neurons (s-LNv), large ventral lateral neurons (l-LNv), dorsal lateral neurons (LNd) and lateral posterior neurons (LPN) (*13*). Recent single cell sequencing has also further divided *Drosophila* clock neurons; there are now 17 different groups without even considering all the “missing” DN3s (*14, 15*) .

Over the last few decades, the functions of many *Drosophila* clock neurons have been extensively studied. For example, adult s-LNvs control morning activity and are essential for free-running rhythms in constant darkness (*16*). In contrast, LNds and non-PDF-expressing s-LNvs modulate both morning and evening activity (*17, 18*). Sleep and activity promoting neurons were both identified within the DN1p population (*19–23*). In contrast, the gene expression, projection patterns and functions of almost all of the 35-40 DN3s – half of the 75 pairs clock neurons -- remain unknown. The singular exception is a recent study suggesting that a small subset of DN3 neurons receive signals from DN1 circadian neurons and then project to “claw” neurons to promote sleep (*23, 24*).

In this study, we developed a novel split GAL4 driver line that is expressed in almost all of the DN3 neurons. To learn more about their gene expression profiles, we utilized a commercially available single cell RNA sequencing method and generated six time points of single cell data around the clock. The 40 DN3 neurons were separated into 12 different clusters by an unsupervised clustering algorithm. Interestingly, some DN3 clusters had striking and unprecedented core clock gene expression patterns; these clusters have poor clock gene transcript cycling as well as a very limited number of cycling transcripts. Moreover, we show that different subsets of DN3 neurons regulate the sleep behavior of flies in a time-of-day dependent manner.

## Results

### A complete high-resolution circadian network transcriptomic cell type atlas

To profile previously uncharacterized DN3 clock neurons, we screened new split GAL4 driver lines. Briefly, carefully chosen activation domains (AD) and DNA binding domains (DBD), driven by two different regulatory regions, were combined so that a functional GAL4 protein was reconstituted only in the overlapping expression regions. The split GAL4 system offers greater specificity compared to the regular GAL4 lines (*25*). We found that the combination of R11B03-AD, from *clockwork orange* (*cwo*), and VT002670-DBD, from vrille (*vri*), labels all or nearly all of the 40 DN3 clock neurons. It also labels the two DN1a, seven DN1p neurons and the s-LNvs with very little ectopic expression (Fig. S1A). To our knowledge, this is the most comprehensive DN3 driver developed to date.

We next combined the R11B03-AD; VT002670-DBD with Clk856-GAL4, which has been used in our previous single cell RNA sequencing studies (*14, 15*) . The combined driver line was then used to drive a nucleus localized EGFP expression. Importantly, there were very few central brain ectopic cells as most of the GFP-expressing neurons were also TIM-positive (Fig. 1A). This strategy allowed us to investigate universal clock neuron gene expression including the DN3s. Moreover, it enabled us to compare DN3 gene expression to previously characterized clock neurons side by side with very limited batch effect issues.

**Figure 1.**
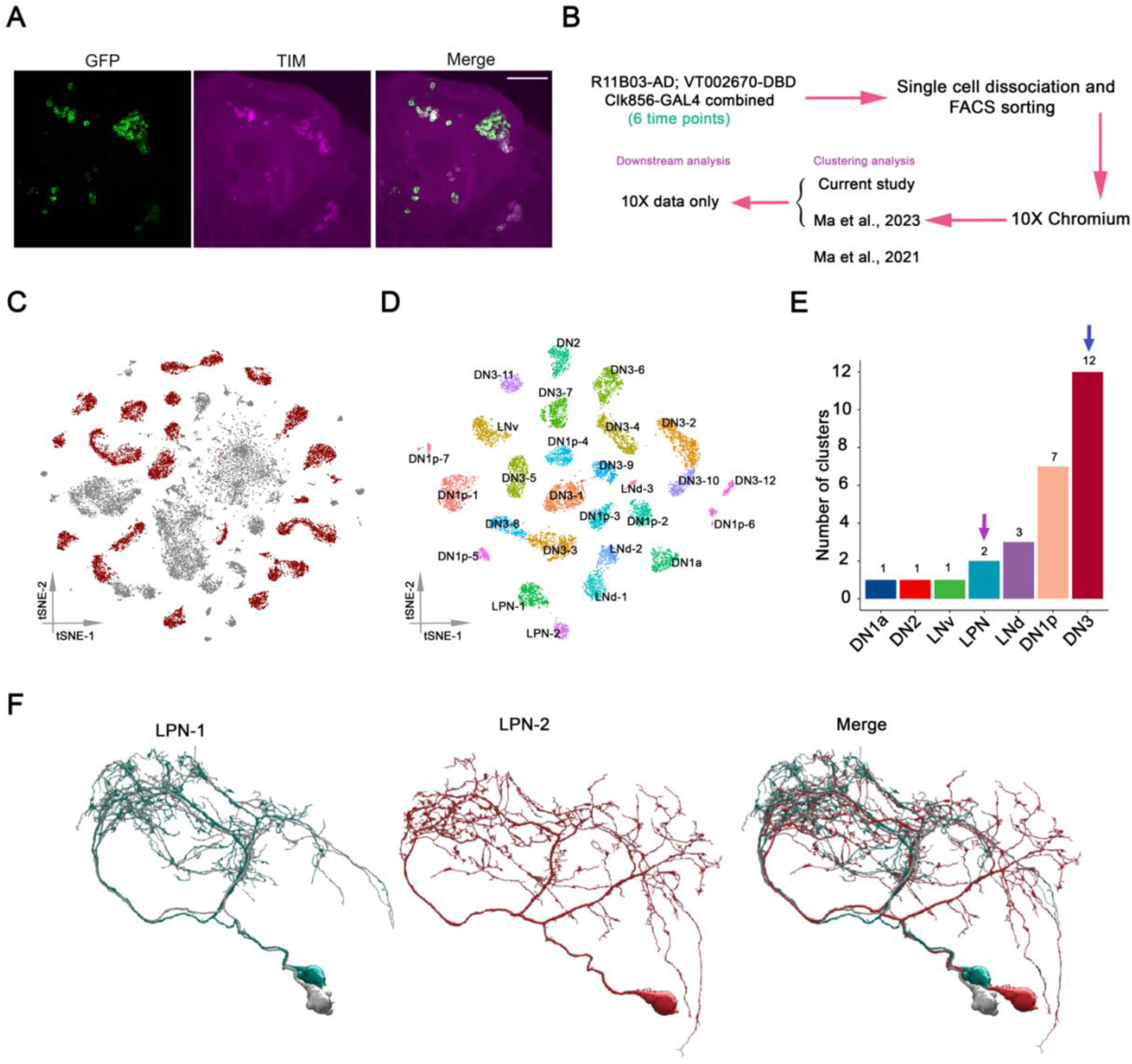
Transcriptomic taxonomy of all *Drosophila* clock neurons. (A) Confocal stack images of immunostained brains from Clk856-GAL4, R11B03-AD; VT2670-DBD > UAS-Stinger-GFP flies at ZT18. Anti-GFP (left), anti-TIM (middle) and a merge of these two images (right). The scale bar represents 50 μm. (B) Schematic workflow of single-cell RNA sequencing around the clock. R11B03-AD; VT2670-DBD was crossed with a stable line of Clk856-GAL4 > UAS-EGFP, the flies were entrained for at least 3 days in LD condition before dissection. The single cell RNA sequencing libraries were generated by 10X Chromium. High confidence single cells were co-clustered with the clock neurons identified previously. (C) t-distributed stochastic neighbor embedding (t-SNE) plot showing the 26210 cells grouped into 69 clusters. High-confidence clusters are shown in purple. (C) t-SNE plot of 10388 *Drosophila* clock in 27 high-confidence clock clusters. The clusters are colored by their cell types. (E) The number of transcriptomic cell types from different anatomic locations. The arrows indicate the LPN and DN3 clock neurons, respectively. (F) Electron microscopy (EM) reconstructions of three LPN clock neurons from the hemibrain data set.

To profile these clock neurons, flies were entrained in a 12:12 Light:Dark (LD) cycle conditions for at least 3 days before they were collected at different times of day. Dissected brains were dissociated by papain and triturated to achieve single cell suspensions before being subject to fluorescence-activated cell sorting (FACS). The single cell RNA sequencing libraries were generated by a commercially available high throughput droplet-based method (10X Chromium).

At each time point, between 2631 to 6913 single cells were assayed (Fig. S1B). To focus on the high confidence single cells, we first applied a cell-wise filtering with the following cycling cutoffs: the number of detected transcripts was between 1000 to 25000, the number of detected genes between 300 to 3500, an entropy of gene expression greater than 5, and a percentage of mitochondrial RNA of less than 10%. After this stringent filtering, we had 9459 high confidence single cells for our clustering analysis. We next combined these high confidence single cells with previously characterized clock neurons, which were similarly subjected to LD entrainment (Fig. 1B and Fig. S1C). The combined dataset not only gave us more statistical power, but the previously identified clock neuron clusters also served as a benchmark for the new clustering.

The average number of detected genes and transcripts in our combined dataset were 1799 and 7304, respectively (Fig. S1D). An unsupervised clustering algorithm was used to integrate the single cell data from the different time points (*26*), resulting in 69 distinct clusters (Fig. 1C). A subsequent cluster-wide filtering based on core clock gene expression resulted in a final total of 27 high confidence clock neuron clusters. Importantly, single cells from our earlier CEL-seq2 data and these 10X data were co-clustered (Fig. S1E-F).

To assign cell identities to the different clusters, we compared previously known single cell data with cells from current study. Interestingly, many clusters show a high degree of correspondence with previously known single cells, suggesting that these clusters were highly reproducible in our current study (Fig. S1G); the Trissin-expressing LNds were an exception (see Discussion). Based on this information, we assigned cell identities to all 27 clusters (Fig. 1D). There are 10 new clusters: they only contain cells only from this current study and reflect DN3 neurons never previously characterized. Combined with the 2 DN3 clusters previously identified, this makes a total of 12 DN3 clusters and 15 other clock neurons clusters (Fig. 1E).

To rule out possible complications from combining more limited CEL-Seq2 data and the 10X data containing the 10 new DN3 clusters, we focused only on the 10X data for subsequent analyses. There are cells from 6 time points in all these clusters (Fig. S2). There is also a unique combination of marker gene expression associated with each cluster (Fig. S3A), and previously known cell type specific marker genes, for example *Pdf*, *ITP*, *CCHa1*, are all expressed in a cell type-specific manner (Fig. S3B). This further raise confidence in the clustering assignments.

There are three LPN neurons in each hemisphere of the fly brain (*27*). Although they were previously in a single cluster (*14, 15*), the current single cell data separated them into two different clusters. This molecular characterization is strikingly consistent with the EM dataset and the morphology described by Reinhard et al., (Fig. 1F): the two sleep-promoting cells in LPN_1 cluster have very similar projections; the wakefulness-promoting LPN_2 has a different pattern characterized by bifurcated branches (*23, 28–30*).

### Mapping DN3 clock neuron gene expression subtypes

To examine gene expression similarity between clock neuron clusters, we carried out gene expression correlation analysis. The LNd, LNv and two DN1p neuron clusters show higher gene expression correlation (Fig. 2A, bottom right), in agreement with a previous result (*14*). Notably, most of the DN3 neuron clusters also have very similar gene expression (Fig. 2A, top left); the exceptions are DN3_7, DN3_11 and DN3_12 (Fig. 2A, purple arrows). As noted above, DN3_7 and DN3_11 were identified previously, indicating that they represent the DN3 neurons which are also labeled by Clk856-GAL4.

**Figure 2.**
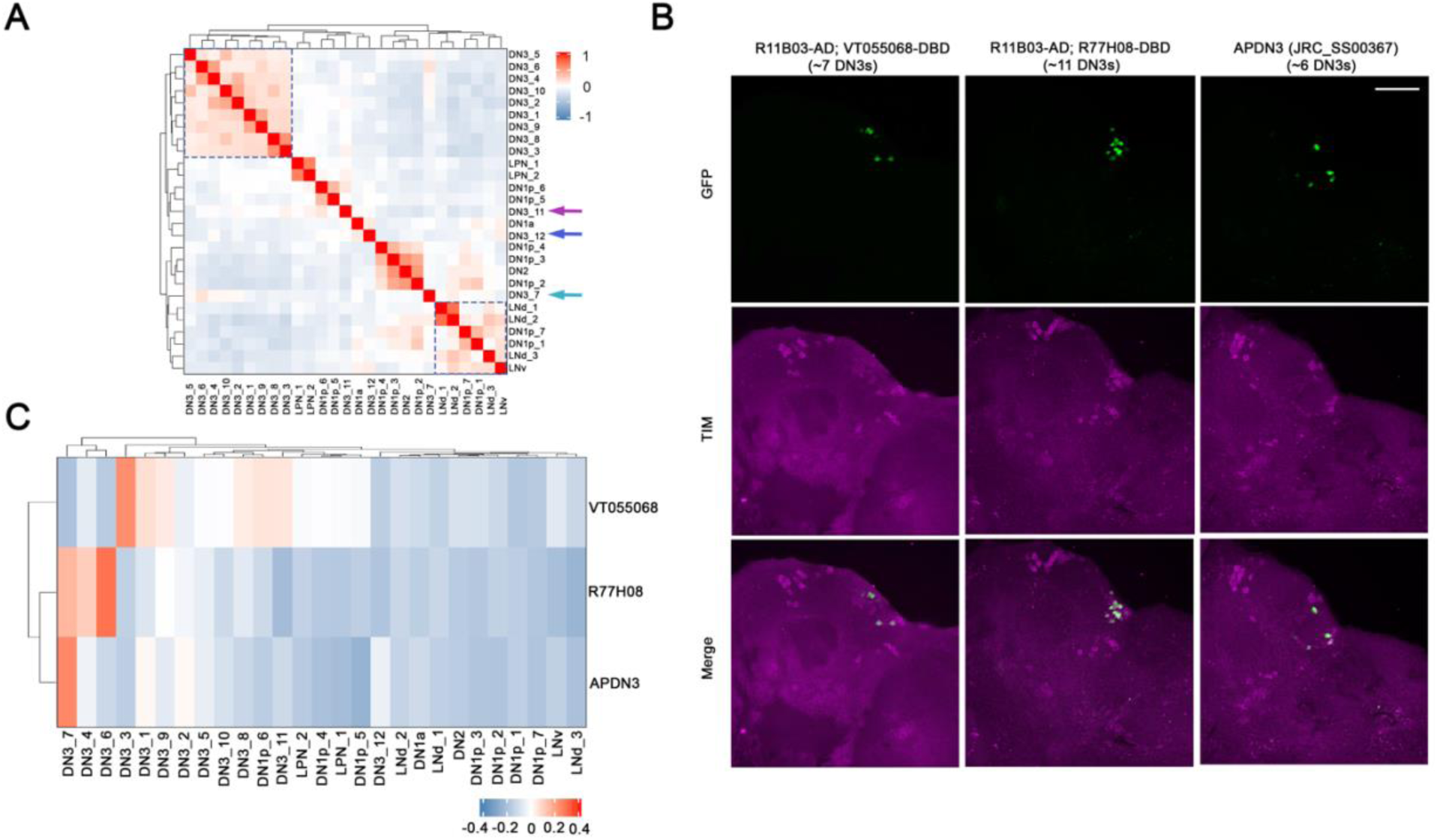
Mapping single cell clusters to DN3 clock neurons. (A) Gene expression correlation of clock neuron clusters. Many of them show higher gene expression correlation except the 3 DN3 clusters pointed by colored arrows. (B) Expression pattern of sparse DN3 split GAL4 drivers. The driver lines were crossed with UAS-Stinger-GFP, flies were entrained in LD for 3 days before dissection at ZT18. Fly brains were co-stained with anti-GFP (green) and TIM (magenta). Scale bar is 50 μm. (C) Gene expression correlation between single cell clusters and DN3 subgroups labeled by different sparse drivers. Flies were entrained in LD for 3 days and collected at ZT02.

To further map the DN3 single cell clusters to brain neurons, we identified 3 sparser DN3-relevant split GAL4s. R11B03-AD; R42F08-DBD is expressed in about 27 DN3 neurons but also labels some DN1s, s-LNvs and a few ectopic cells. The other two split GAL4s, R11B03-AD; VT055068 and R11B03-AD; R77H08, label very specifically only 7 and 11 DN3 neurons, respectively (Fig. 2B). To assess the gene expression profiles of specific DN3 neurons, these new DN3 driver lines as well as driver SS00367, which labels sleep-promoting APDN3 neurons (*24*), were used to drive EGFP expression. Cells were collected at ZT02, RNA sequencing libraries were generated with Smart-seq3 (*31*), and these bulk RNA seq data compared to the single cell data (Fig. 2C). The analysis indicates that the APDN3s are most highly correlated with a single cluster, DN3_7. The DN3 neurons identified by R11B03-AD; R77H08 correspond to 3 clusters: DN3_7 and therefore also to APDN3s as well as to cluster DN_4 and more strongly to cluster DN3_6. R11B03-AD; VT055068-DBD was most highly correlated with DN3_3 and more weekly with several other clusters (Fig. 2C).

### Unprecedented rhythmic gene expression in DN3s

We suspected that DN3 clock neuron clusters have rhythmic gene expression patterns like the other clock neurons clusters already characterized (*14*). Indeed, the clock gene *vri* shows a canonical temporal pattern in most of the DN3 clusters with peak expression between ZT14 to ZT18 (Fig. 3A). However, 3 DN3 clusters have unprecedented *vri* patterns: the cycling amplitude of DN3_3, DN3_4 and DN3_8 is much reduced with substantial expression at trough times, early morning and light night; moreover, the very modest peak of *vri* expression in DN3_3 is delayed relative to the peak time of other clusters (Fig 3A blue arrows). The *pdp1* cycling patterns in these 3 exceptional clusters are similar (Fig. S4, blue arrows). The *tim* cycling patterns in 2 of these clusters, DN3_3 and DN3_8, are even more unusual than their *vri* and *pdp1* patterns; these 2 *tim* patterns show exceptionally high trough expression levels. Moreover, the delayed *tim* expression peak in DN3_3 is even more apparent than its delayed *vri* expression peak (Fig 3B), which indicates that delayed clock gene expression in DN3_3 is probably general.

**Figure 3.**
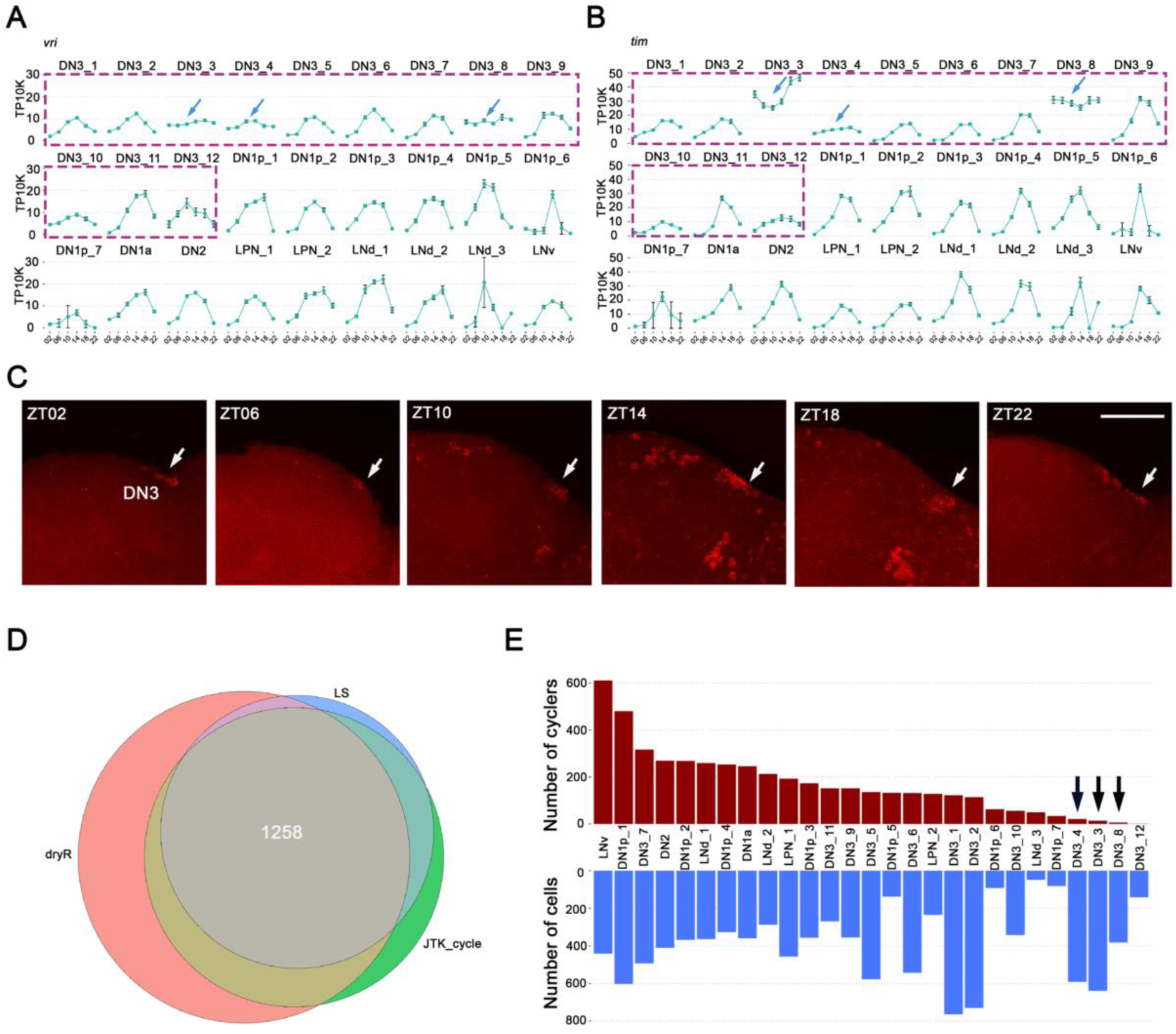
Unprecedented *tim* expression in DN3 neurons. (A-B) The mean *vri* (A) and *tim* (B) expression throughout the day in light: dark (LD) condition is graphed for each cluster. Error bars represent mean ± SEM. The arrows indicate the three DN3 clusters showing shifted or dampened *vri* or *tim* expression. (C) Fluorescent in situ hybridization (FISH) for *tim* mRNA transcripts in DN3 neurons throughout the day in LD condition. (D) Venn diagram showing the identified cycling transcripts by JTK_Cycle, LS and dryR. There are 1258 high confidence cycling transcripts in total. (E) The number of cells (bottom) and cycling transcripts in each cluster (top). The black arrows indicate the three DN3 clusters showing very limited rhythmic gene expression.

To further verify the unusual clock gene expression patterns in DN3 neurons, we carried out single molecular Fluorescence in situ hybridization (FISH) in cleared whole-mount adult *Drosophila* brains (*32*). *tim* mRNA is clearly detectable only in the DN3s at traditional trough times of day (Fig 3C), which further underscores the have unusual clock gene expression features of some DN3 neurons.

To address rhythmic gene expression of DN3s more generally, we used three standard algorithms to define oscillating gene expression among the clock neuron clusters: JTK_Cycle, Lomb-Scargle (LS) and dryR (*33–35*). The following cycling cutoffs were used to focus on robust cycling transcripts: a JTK_cycle p-value of less than 0.05, a LS p-value of less than 0.05, a dryR p-value of less than 0.0, a cycling amplitude (maximum expression divided by minimum expression) of at least 2-fold, and a maximal expression of at least 0.5 TP10K. In total, 1258 cycling transcripts were identified within the clock neuron clusters by all of these three different methods (Fig. 3D).

The number of cycling transcripts varies greatly between clusters, due in large part to the variable number of recovered cells per cluster. Notably however, DN3_3, DN3_4 and DN3_8 have many fewer cycling transcripts than DN3_7 and DN3_11 despite the fact that the first 3 clusters have more cells than DN3_7 and DN3_11 (Fig. 3E). In light of the the aberrant clock gene cycling profiles expression in DN3_3, DN3_4 and DN3_8, we speculate that the molecular circadian program in these three clusters is less robust than in other circadian neurons clusters (see Discussion).

Neurotransmitters and neuromodulators interact with GPCRs to excite, inhibit or modulate the GPCR-expressing neurons. Our recent studies have shown that GPCRs are highly enriched in the fly brain clock network, and GPCR transcripts alone can define the adult brain dopaminergic and circadian neurons (*15, 36*). Notable in this context, *CapaR* encodes a GPCR for the neuropeptide Capa, and this transcript is substantially and rhythmically expressed in only a single DN3 cluster, DN3_11 (Fig. S5). A second GPCR transcript, *TrissinR,* is highly expressed in 7 different DN3 clusters (Fig. 4A) and oscillates robustly in clusters DN3_5, DN3_6 and DN3_7 (Fig. 4B). Its ligand, the neuropeptide Trissin, is expressed in only two LNd clock neurons as well as a few neurons outside of the clock network (*14, 37*).

**Figure 4.**
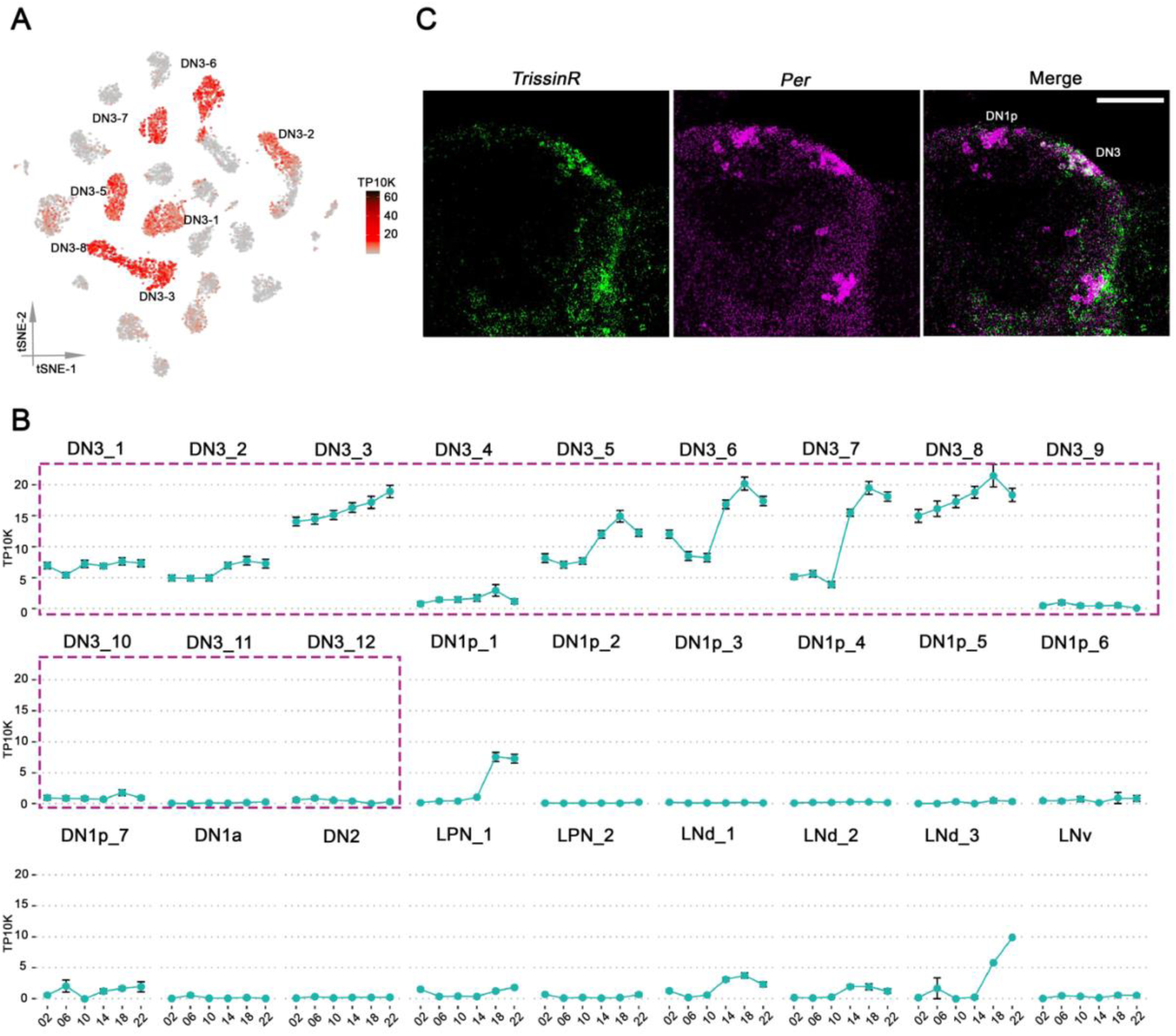
*TrissinR* expression in Drosophila clock neurons. (A) t-SNE plots showing *TrissinR* expression. Red indicates higher expression (color bar, TP10K). (B) The mean *TrissinR* expression throughout the day in LD condition is graphed for each cluster. Error bars represent mean ± SEM. *TrissinR* expression is specific to the DN3 neurons. (C) RNA-scope for *TrissinR* mRNA transcripts in DN3 neurons at ZT18.

To further verify *TrissinR* expression in DN3s, we conducted in situ hybridization with *TrissinR* probes at ZT18, when TrissinR shows peak expression levels (Fig. 4B). *TrissinR* is indeed enriched in the DN3 region of the fly brain, which is further defined by *per* expression (Fig. 4C).

### Different DN3 subtypes regulate daytime and nighttime sleep

Is there a biological function of TrissinR expression in DN3s? To this end, we exploited our recent *Drosophila* GPCR gRNA library as well as recently characterized split GAL4 driver lines to knock out *TrissinR* only in DN3s. The broad DN3 driver R11B03-AD; VT2670-DBD resulted in a robust reduced sleep phenotype in the daytime as well as the nighttime (Fig. 5A). Knocking down TrissinR expression with the same driver and an RNAi line showed comparable phenotypes (Fig. S6). To further investigate DN3 neuron subsets, we knocked out *TrissinR* only in the approximately 11 DN3 neurons expressed by the R11B03-AD; R77H08-DBD driver. These flies only showed reduced nighttime sleep (Fig. 5B). Given the much more modest effect of knocking out *TrissinR* with the APDN3 driver and the expression patterns of the R11B03-AD; R77H08-DBD driver (Fig. 2C), we suggest that much of the *TrissinR* nighttime sleep-promoting effect comes from DN3 cluster 6. We further speculate that DN3s and their TrissinR expression have a general sleep-promoting role and that distinct DN3 subgroups regulate different features of sleep by interacting with different downstream targets in a time-dependent manner (see Discussion).

**Figure 5.**
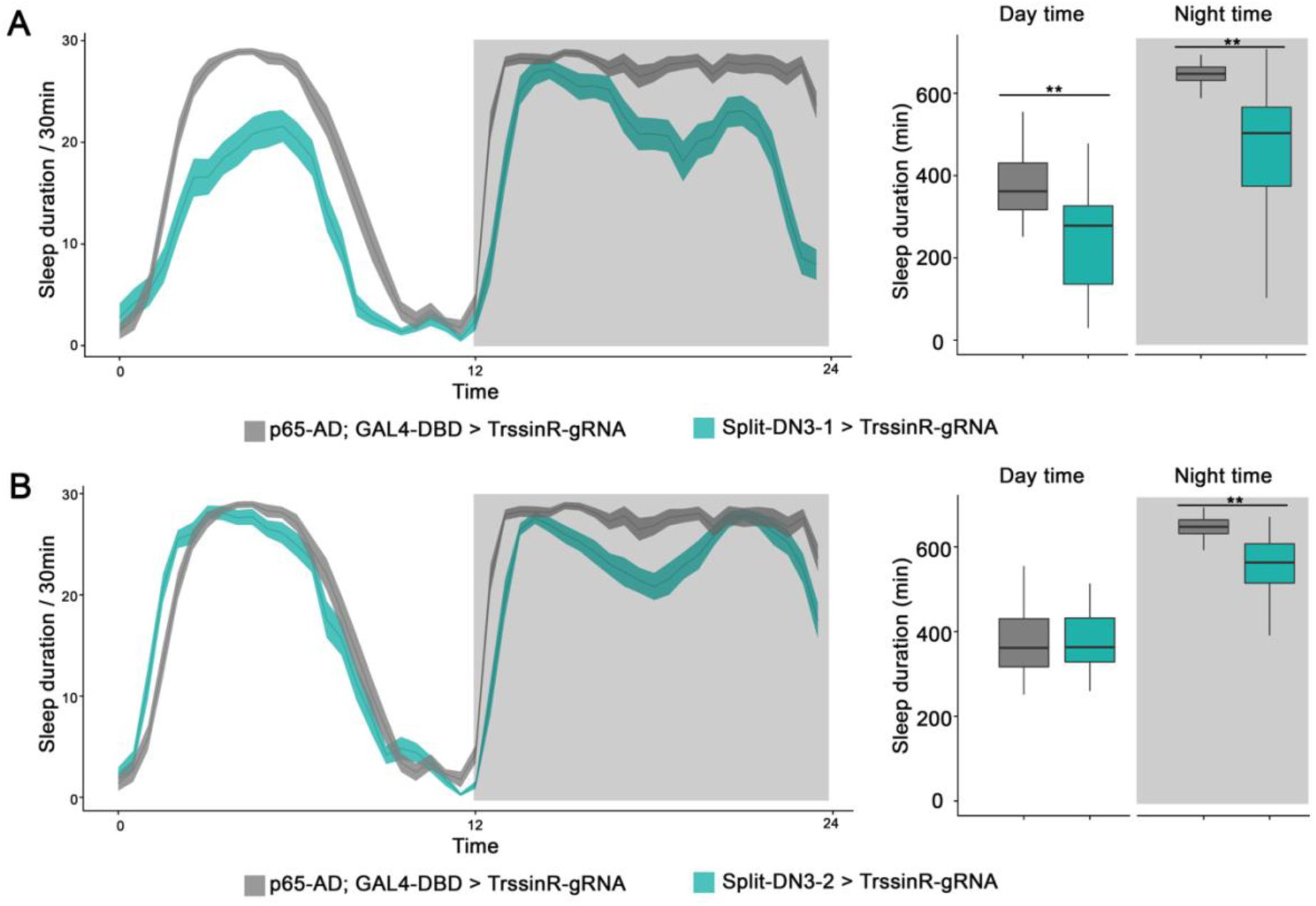
TrissinR expression in DN3 neurons promotes sleep at different times of day. (A-B) Sleep plots of R11B03-AD; VT2670-DBD (A), R11B03-AD; R77H08-DBD (B) driven UAS-TrissinR-gRNA; UAS-Cas9 (blue) and controls (gray). The solid lines represent the averaged sleep amount, and the shading represents SEM for each time point. Quantified daytime and nighttime sleep durations are shown on the right panels.

## Discussion

This study provides an invaluable resource, namely, the most comprehensive and high-resolution single cell transcriptomic analysis of the 150 *Drosophila* clock neurons with a particular focus on the previously uncharacterized DN3s. This medium size number of clock network neurons makes it an ideal system to study how gene expression patterns within different transcriptomic cell types contribute to distinct projection patterns and ultimately a variety of circadian behaviors. Indeed, gene expression very likely underlies the heterogeneous morphologies of different clock neurons. Moreover, different DN3 subtypes regulate the sleep behavior of flies in a time dependent manner.

It is perhaps surprising that the about 150 clock neurons in Drosophila brain are separated into so many -- at least 27 – distinct transcriptomic cell types. We previously showed that most of the clock neurons and even dopaminergic neurons in the adult fly brain are heterogeneous at the RNA level (*14, 15*), and this current study extends this conclusion to the enigmatic DN3 neurons. There are about 12 DN3 transcriptomic cell types of which two were previously identified (*14*).

The large number of gene expression cell types still somewhat underrepresents the diversity of gene expression withiin clock neurons. Most of the previous single cell transcriptomic cell types were reproduced in the current study with one exception: it is the Trissin-expressing LNd neurons, which is probably due to the technical bias from the single cell RNA sequencing methods. Nonetheless, the remarkable gene expression diversity described here contrasts with the single cell/nucleus RNA sequencing efforts from mice SCN (*6–8*). In this context, we cannot rule out the possibility that the gene expression heterogeneity in the current study is *Drosophila*-specific considering that clock neurons are established during development (*38*).

In addition to the gene expression classification of clock neurons, our study also allows us to compare the transcriptomic cell types with neuron morphologies and projection patterns. The six LNd clock neurons were separated into 4 different morphological groups, which are in concordance with the EM dataset (*14, 39*). Interestingly, the three LPN neurons are now separated into two different clusters, which is likely due to the good cell coverage provided by the current study and reflects the high fidelity of our single cell gene expression analysis. It is of great interest how these two adjacent LPN groups interact with their partners to regulate sleep and wakefulness, respectively.

Most of the lateral clock neurons and some of the dorsal clock neurons have been extensively studied, but the gene expression basis and functions of DN3 clock neurons are largely unknown. One main reason for this deficiency is the lack of good driver lines to label these neurons. The DN3 neurons appear to be heterogeneous based on previous immunohistochemistry studies (*29, 40, 41*). The joint effort of split GAL4 screening and single cell RNA sequencing in the current study resolved the gene expression basis of DN3 clock neuron identity for the first time in decades. The methodology in the current study is readily applicable to other neurons in *Drosophila* adult brain.

Rhythmic gene expression in the brain and peripheral tissues dictates many different behaviors and at least 80% of protein coding genes are reported to be oscillating in a daily manner in a tissue-specific fashion (*42, 43*). In this context, some DN3 clusters had unusual cycling features, i.e., their core clock gene expression was shifted or showed a dampened amplitude. More general rhythmic gene expression analysis also indicated that there were very few cycling transcripts in these clusters, especially in DN3_3, DN3_4 and DN3_8. We noted the the two DN3 clusters labeled by Clk856-GAL4 showed more robust core clock gene cycling as well as cycling transcript expression in general. How core clock gene expression is differentially regulated in these neurons needs further investigation.

GPCRs play a pivotal role in neuron communication. Our previous studies showed that GPCRs are enriched in *Drosophila* clock neurons, and GPCRs alone could define the cell types within the clock network (*15, 36*). Indeed, a large number of GPCRs are rhythmically expressed in a cell type specific manner, which further underscores the importance of the dynamic interactions between the circadian network and their upstream or pre-synaptic partner neurons. Additionally, our analysis of cell type specific GPCR expression provides novel functional insight into DN3 function. We show that TrissinR expression in different DN3 subtypes promote fly sleep in a time dependent manner. These new findings complement our previous study showing that a small subset of DN3 neurons receive synaptic input from DN1ps and project to claw neurons to promote fly sleep (*24*).

In summary, the DN3 clock neurons in *Drosophila* adult brain are strikingly heterogeneous in gene expression and function. Moreover, the newly developed driver lines and many cell type specific marker genes identified in this study will help drive further understanding of DN3 clock neuron function.

## Materials and methods

### Fly strains and rearing

Flies were raised on standard cornmeal medium with yeast under LD (12 h light/12 h dark) conditions at room temperature. The fly lines used in this study are listed in the key resource table.

### Fluorescence-activated cell sorting

To label the clock neurons, we first generated a stable line of Clk856-GAL4; UAS-EGFP, and then crossed it with R11B03-AD; VT2670-DBD. Equal numbers of males and females were first entrained for at least 3 d under LD conditions at 25 °C before dissection at different time points of the day. Fly brains were dissected in cold dissection saline [9.9 mM HepesKOH (pH 7.4), 137 mM NaCl, 5.4 mM KCl, 0.17 mM NaH2PO4, 0.22 mM KH2PO4, 3.3 mM glucose, and 43.8 mM sucrose] with neuronal activity inhibitors (20 μM 6,7-dinitr oquinoxaline-2,3dione, 0.1 μM tetrodotoxin, and 50 μM D, L-2-amino-5-phosphonovaleric acid). The brains were digested with papain (50 U/ml, ∼2 μl per brain; Worthington Biochemical, #LK003176) at room temperature for 25 to 30 min. Brains were then washed twice with ice-cold active Schneider’s Medium (SM) medium after the digestion. Flame-rounded 1000-μl pipette tips with different sized openings were used to triturate the brains. The resulting cell suspension was passed through a 100-μm sieve. Hoechst dye (one drop per 0.5 ml of sample; Invitrogen, #R37605) was added into the sample tube to stain the nucleus (15 minutes at room temperature). A BD Melody fluorescence-activated cell sorting (FACS) machine in single-cell sorting mode was used for cell collection. GFP- and Hoechst-positive single cells were collected in a 1.5-ml Eppendorf tube containing 0.3 ml of collection buffer [phosphate-buffered saline (PBS) + 0.04% bovine serum albumin] and used for downstream single cell RNA sequencing library preparation. To minimize the possible stress to the cells, the collection devices were kept at 5°C constantly during the sorting process.

### Library preparation and raw data processing

The collected cells from the FACS machine were spun down on a benchtop centrifuge by 700g for 10 min at 4°C before loaded to the GEM chip from Chromium Single Cell 3′ Kit (v3) of 10X Genomics. The libraries were prepared according to the standard user guide (CG000315 Rev. B) from 10X without any modifications. 10X libraries were sequenced by Illumina NextSeq 500 with the High Output Kit v2.5 (75 cycles). The Cell Ranger package from 10X was used to map the sequencing data to the *Drosophila* genome (dm6), only the alignments to annotated exons were used for UMI quantitation.

### Dimensionality reduction and clustering analysis

To focus on the high confident single cells, we first filter out low-quality cells based on the following criteria: (i) fewer than 300 or more than 3500 detected genes; (ii) fewer than 1000 or more than 15000 total UMI; (iii) gene expression entropy smaller than 5.0, where entropy was defined as −nUMI x ln(nUMI) for genes with nUMI > 0, where nUMI was a number of UMI in a cell; and (iv) the percentage of mitochondria RNA greater than ten percent. The possible doublets were detected by Scrublet with default setting, and excluded from the downstream analysis.

We next combined the high confidence clock neurons from previous studies and the single cells from current study to identify the facile clock neuron clusters (*14, 15*). The integration and clustering method has been described previously (*9*). We first separate the raw dataset by methods and time points, then carry out data transformation, normalization and variance by SCTransform function from the Seurat package (*26*). The batch effect was removed by regressing out numbers of genes, UMIs, detected genes per cell, sequencing batches, and percentage of mitochondrial transcripts. We computed 3000 variable genes at each time point and method and only used the shared variable genes in all conditions for the clustering analysis, which resulted in 69 clusters. We next applied a cluster wise filtering, only the clusters with high core clock genes are retained.

### Bulk RNA sequencing

Flies were entrained for at least 3 days before collected and dissected at ZT02. GFP positive neurons were sorted and collected as aforementioned. PolyA mRNA was isolated with the Dynabeads mRNA direct kit (Thermo fisher 61011). Subsequently, complementary DNA (cDNA) and sequencing libraries were prepared using Smart-seq3 (*31*). Final libraries were quantified on a High Sensitivity D1000 ScreenTape on the Agilent TapeStation.

### Cycling transcripts analysis

We used three different computational methods, including JTK_Cycle, LS and dryR. To identify the facile cycling transcripts in single cell clusters (*33–35*). To be considered cycling, the following cutoffs were used: (i) a JTK_cycle p-value of less than 0.05; (ii) a LS p-value of less than 0.05; (iii) a dryR p-value of less than 0.01; (iv) a cycling amplitude (maximum expression divided by minimum expression) of at least 2-fold, and (v) a maximal expression of at least 0.5 TP10K.

### Immunostaining

Three to five days old flies were entrained in LD conditions for 3 days before being fixed with 4% (vol/vol) paraformaldehyde with 0.5% Triton X-100 for 2 hr and 40 min at room temperature. Brains were dissected and then washed twice (10 min) in 0.5% PBST buffer before being blocked overnight in 10% Normal Goat Serum (NGS; Jackson ImmunoResearch Lab) at 4 °C. The brains were then incubated in, a rat anti-TIM at 1:200 dilution, a mouse or chicken anti-GFP antibody at a 1:1000, a rat anti-RFP antibody at 1:200 for overnight, then the brains then washed twice (10 min) in 0.5% PBST buffer. The corresponding secondary antibodies were added and incubated overnight. Brains were mounted in Vectashield (Thermal Fisher) and imaged on a Leica SP8 confocal microscope.

### Fluorescent in situ hybridization

Fluorescent in situ hybridization (FISH) for *timeless* expression was performed as described previously onto wild-type flies (w1118) at different times of day with the following exceptions: custom oligo probes were ordered against the entire *timeless* mRNA sequence and conjugated with Quasar 670 dye (Stellaris Probes, Biosearch Technologies). The probes were diluted to a final concentration of 0.75 mM for the hybridization reaction. The brains were washed before mounting onto slides with Vectashield Mounting Medium (Vector Laboratories). The slides were immediately viewed on a Zeiss 880 series confocal microscope.

### Drosophila behavior assay

We used the *Drosophila* Activity Monitoring system (Trikinetics) to record the number of beam crosses caused by the fly in one-min intervals. Five to seven-day-old male flies were used for the behavior analysis, in which flies were individually placed in glass tubes containing 2% agar and 4% sucrose. The flies were entrained under 12:12 LD conditions for at least 3 days. Each experiment was performed at least twice and got similar results. The sleep analysis was performed with MATLAB. Statistical analysis was performed using a one-way analysis of variance (ANOVA) (www.statskingdom.com), and a p value of less than 0.05 compared to all control groups was considered significant.

## Acknowledgements

We thank the members of Rosbash lab for critical reading and thoughtful discussion of the manuscript. We greatly appreciate Drs. Fang Guo and Yerbol Kurmangaliyev for insightful comments on the manuscript. Funding: This work was supported by Howard Hughes Medical Institute. Author conctributions: D.M, K.Z and M.R designed the study. D.M, J.Q.L, X.D and M.M.D prerfomed the experiments and data analysis. D.M, K.C.A and M.R wrote the paper.

---

**Figure S1.**
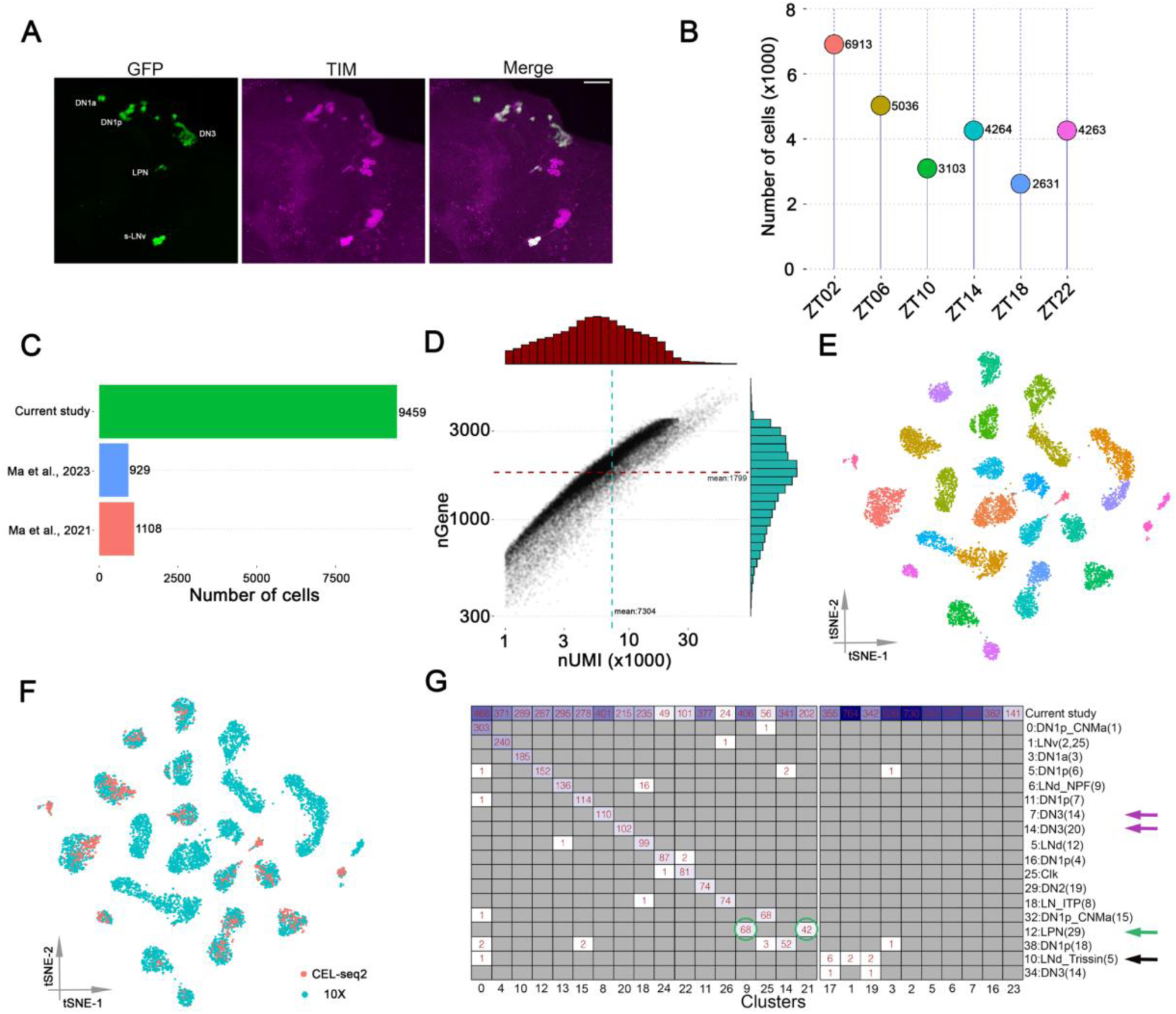
Identification of high confidence *Drosophila* clock neuron clusters. (A) Confocal stack images of immunostained brains from R11B03-AD; VT2670-DBD > UAS-Stinger-GFP flies at ZT18. Anti-GFP (left), anti-TIM (middle) and a merge of these two images (right). The scale bar represents 50 μm. (B) The number of single cells from 10X Chromium at each time point before initial filtering. (C) The number of high confidence single cells for the clustering analysis. (D) The mean detected genes and transcripts in the combined dataset. (E) t-SNE plot showing the 27 high-confidence clock clusters after the cluster wise filtering. The clusters are colored by their cell types. (F) CEL-Seq2 data and 10X data are co-clustered in different clusters. CEL-Seq2 data are shown in red and 10X data are shown in blue. Two previously identified DN3 clusters are highlighted in circles. (G) Heatmap showing the high degree of correspondence of the single cells from previous and current studies. The arrows in magenta indicate two previously identified DN3 clusters, the arrow and circles in blue indicate the two LPN clusters, and the arrow in black indicate the Trissin-expression LNds.

**Figure S2.**
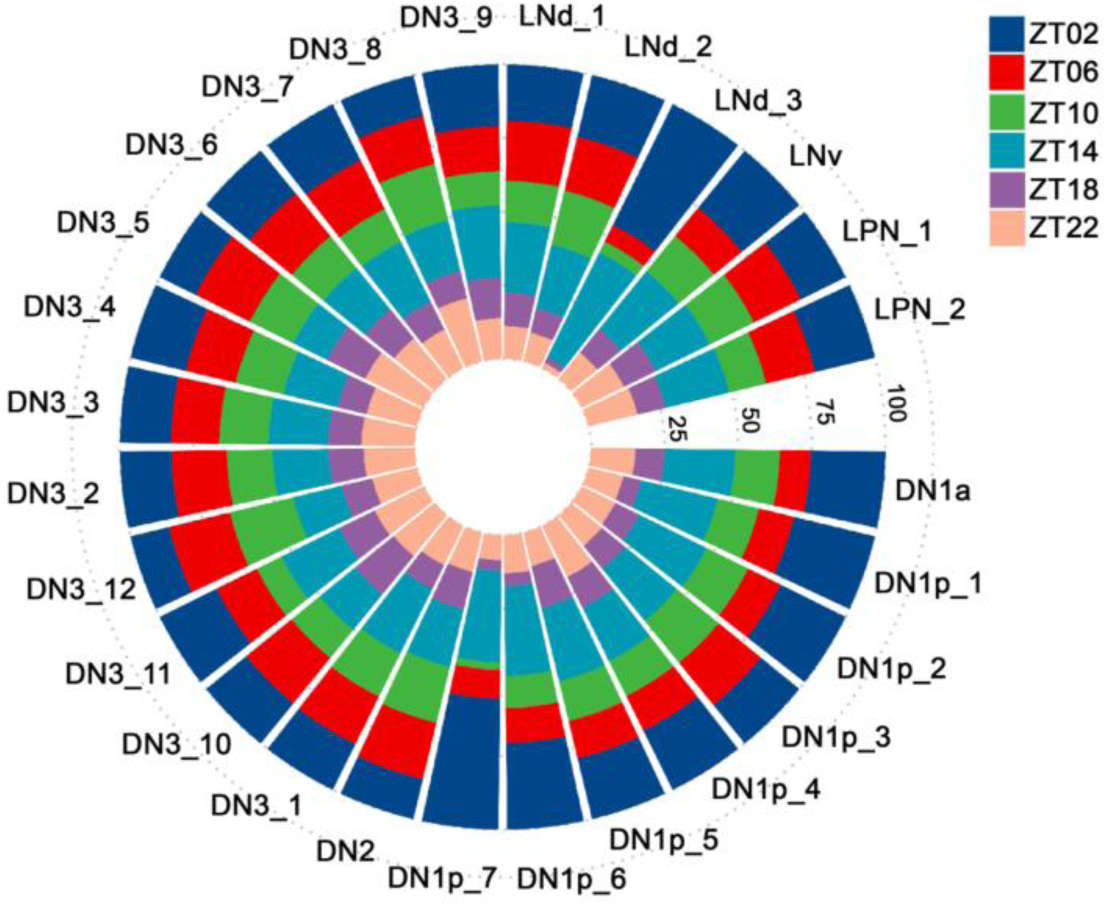
The percentage of cells from six time points in each single cell cluster. Circled bar plot showing that in high confidence clusters there are cells from 6 time points in Light: Dark conditions. Each time point is represented by different colors.

**Figure S3.**
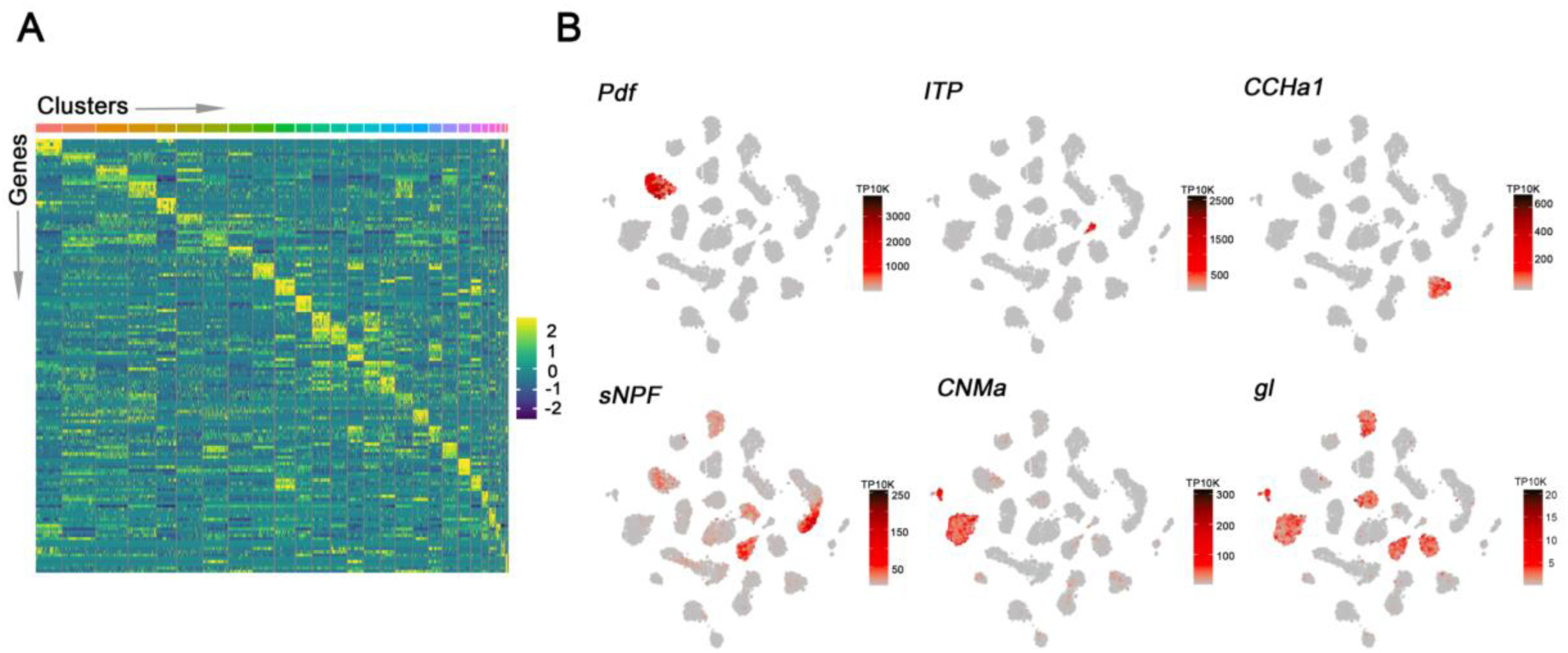
Marker gene expression in each cluster. (A) Heatmap showing the expression levels of the top 5 differentially expressed genes (rows) in cells (columns). Clusters are ordered by size and are represented by different colors on top of the heatmap. (B) t-SNE plots showing previously known marker gene expression in all clusters. Each cell is colored by the expression levels with gray indicating low expression and black indicating the highest expression.

**Figure S4.**
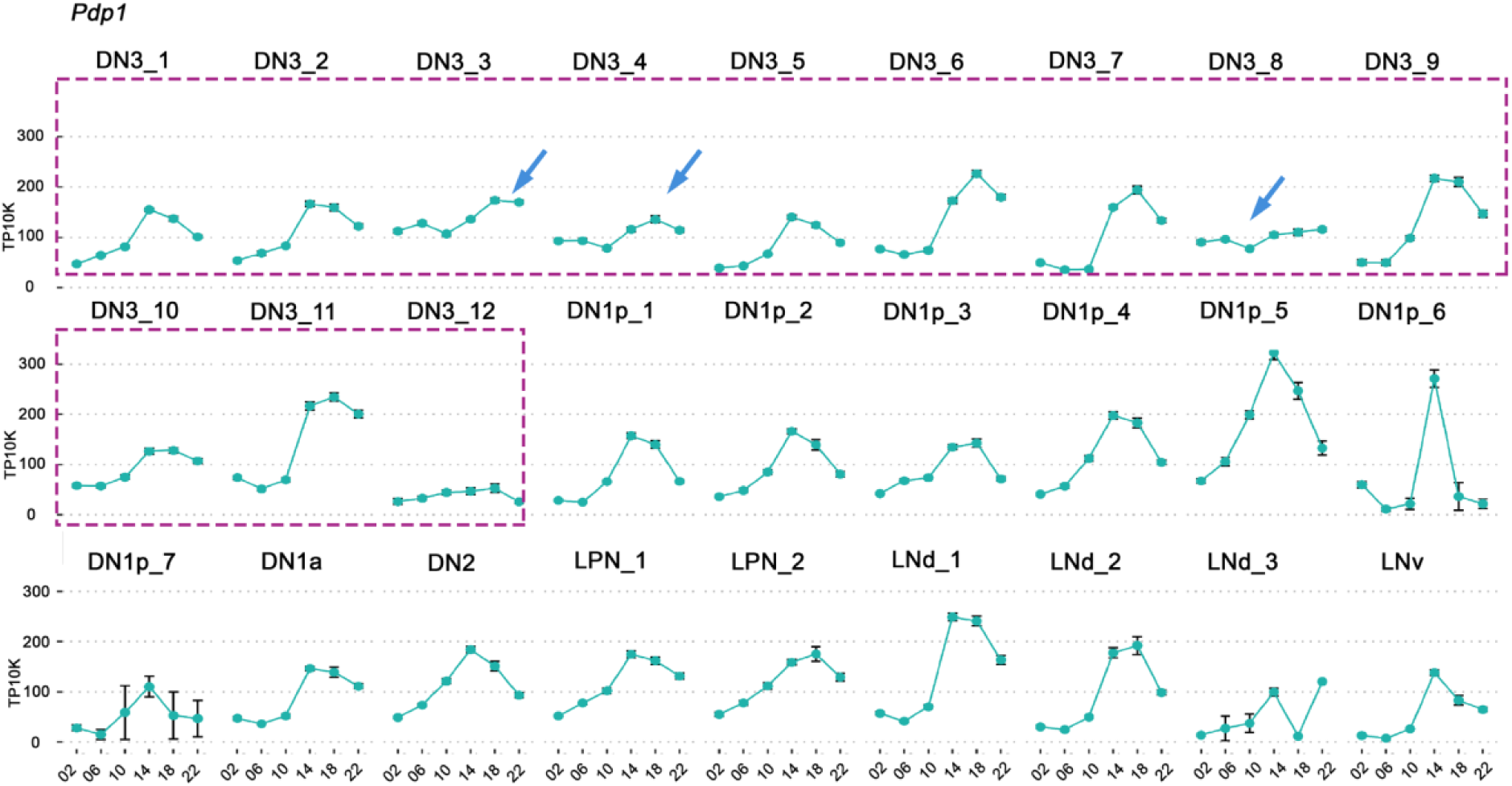
*Pdp1* expression in each cluster. The mean *Pdp1* expression throughout the day in LD conditionis is graphed for each cluster. Error bars represent mean ± SEM. The arrows indicate the three DN3 clusters showing shifted or dampened *Pdp1* expression.

**Figure S5.**
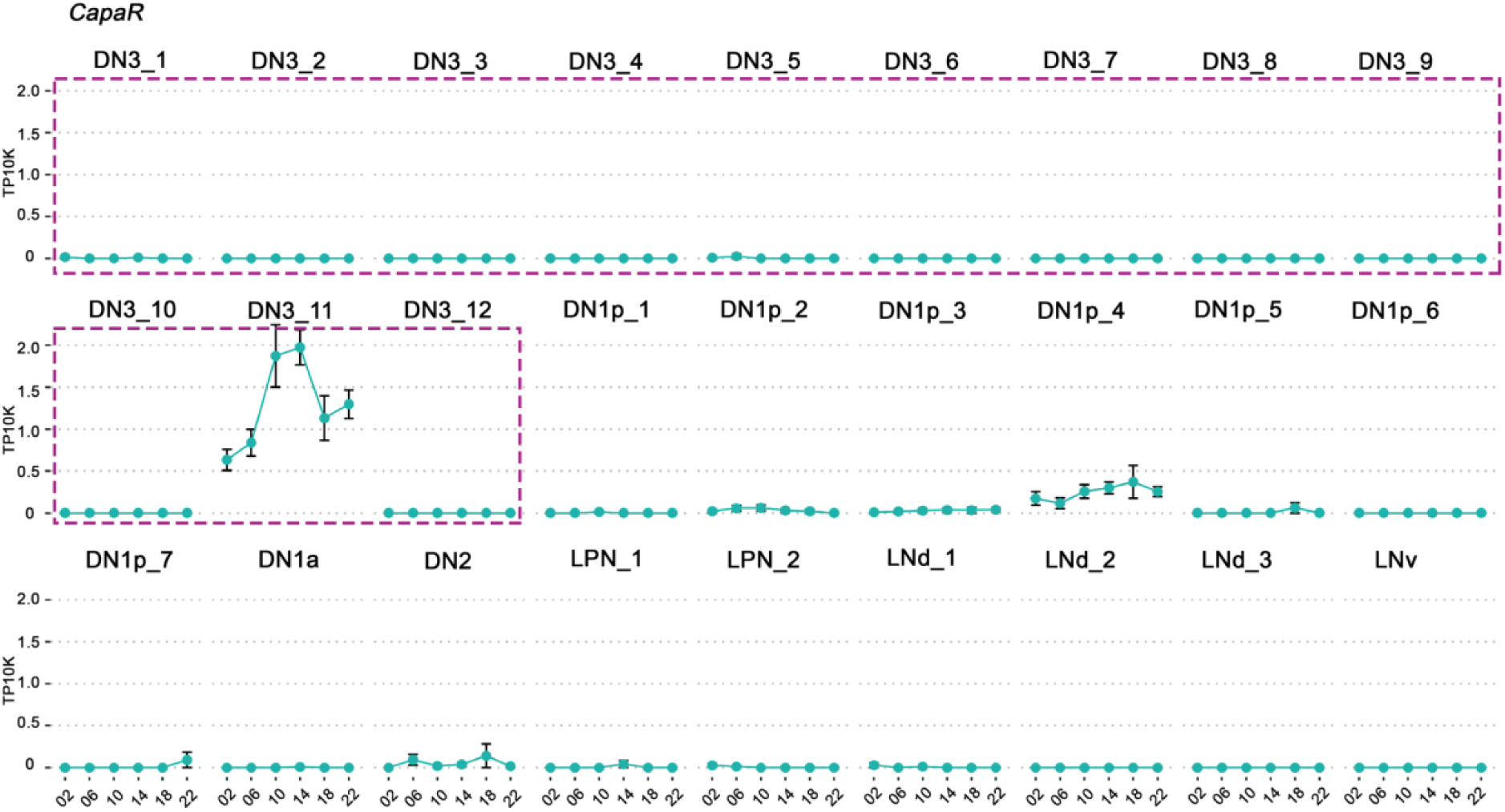
*CapaR* expression in each cluster. The mean *CapaR* expression throughout the day in LD conditionis is graphed for each cluster. Error bars represent mean ± SEM. The arrows indicate the three DN3 clusters showing shifted or dampened *CapaR* expression.

**Figure S6.**
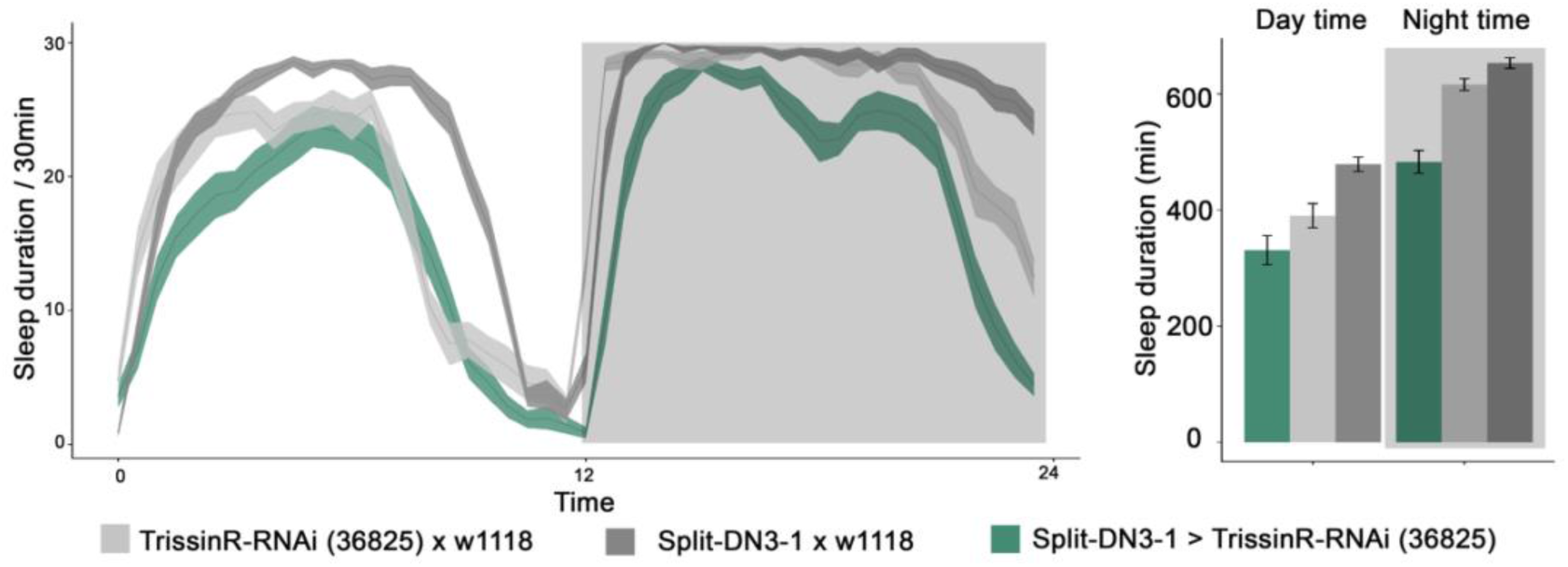
TrissinR expression in DN3 neurons promotes sleep at different times of day. Sleep plots of R11B03-AD; VT2670-DBD driven UAS-TrissinR-RNAi (36825) and controls (gray). The solid lines represent the averaged sleep amount, and the shading represents SEM for each time point. Quantified daytime and nighttime sleep durations are shown on the right panel.

## Notes

### Competing Interest Statement

The authors have declared no competing interest.

## References

1. R. Allada, B. Y. Chung, Circadian organization of behavior and physiology in Drosophila. Annual review of physiology 72, 605–624 (2010).

2. S. M. Siepka, S. H. Yoo, J. Park, C. Lee, J. S. Takahashi, Genetics and neurobiology of circadian clocks in mammals. Cold Spring Harb Symp Quant Biol 72, 251–259 (2007).

3. D. Top, M. W. Young, Coordination between Differentially Regulated Circadian Clocks Generates Rhythmic Behavior. Cold Spring Harb Perspect Biol 10, (2018).

4. J. S. Takahashi, Transcriptional architecture of the mammalian circadian clock. Nature reviews. Genetics 18, 164–179 (2017).

5. J. A. Mohawk, J. S. Takahashi, Cell autonomy and synchrony of suprachiasmatic nucleus circadian oscillators. Trends in neurosciences 34, 349–358 (2011).

6. S. Wen, D. Ma, M. Zhao, L. Xie, Q. Wu, L. Gou, C. Zhu, Y. Fan, H. Wang, J. Yan, Spatiotemporal single-cell analysis of gene expression in the mouse suprachiasmatic nucleus. Nature neuroscience 23, 456–467 (2020).

7. P. Xu, S. Berto, A. Kulkarni, B. Jeong, C. Joseph, K. H. Cox, M. E. Greenberg, T. K. Kim, G. Konopka, J. S. Takahashi, NPAS4 regulates the transcriptional response of the suprachiasmatic nucleus to light and circadian behavior. Neuron 109, 3268–3282 e3266 (2021).

8. E. L. Morris, A. P. Patton, J. E. Chesham, A. Crisp, A. Adamson, M. H. Hastings, Single-cell transcriptomics of suprachiasmatic nuclei reveal a Prokineticin-driven circadian network. EMBO J 40, e108614 (2021).

9. J. Lu, Y. H. Zhang, T. C. Chou, S. E. Gaus, J. K. Elmquist, P. Shiromani, C. B. Saper, Contrasting effects of ibotenate lesions of the paraventricular nucleus and subparaventricular zone on sleep-wake cycle and temperature regulation. The Journal of neuroscience : the official journal of the Society for Neuroscience 21, 4864–4874 (2001).

10. S. Deurveilher, K. Semba, Indirect projections from the suprachiasmatic nucleus to major arousal-promoting cell groups in rat: implications for the circadian control of behavioural state. Neuroscience 130, 165–183 (2005).

11. T. C. Chou, T. E. Scammell, J. J. Gooley, S. E. Gaus, C. B. Saper, J. Lu, Critical role of dorsomedial hypothalamic nucleus in a wide range of behavioral circadian rhythms. The Journal of neuroscience : the official journal of the Society for Neuroscience 23, 10691–10702 (2003).

12. N. Vujovic, J. J. Gooley, T. C. Jhou, C. B. Saper, Projections from the subparaventricular zone define four channels of output from the circadian timing system. The Journal of comparative neurology 523, 2714–2737 (2015).

13. C. Dubowy, A. Sehgal, Circadian Rhythms and Sleep in Drosophila melanogaster. Genetics 205, 1373–1397 (2017).

14. D. Ma, D. Przybylski, K. C. Abruzzi, M. Schlichting, Q. Li, X. Long, M. Rosbash, A transcriptomic taxonomy of Drosophila circadian neurons around the clock. eLife 10, (2021).

15. D. Ma, N. Herndon, J. Q. Le, K. C. Abruzzi, K. Zinn, M. Rosbash, Neural connectivity molecules best identify the heterogeneous clock and dopaminergic cell types in the Drosophila adult brain. Sci Adv 9, eade8500 (2023).

16. S. C. Renn, J. H. Park, M. Rosbash, J. C. Hall, P. H. Taghert, A pdf neuropeptide gene mutation and ablation of PDF neurons each cause severe abnormalities of behavioral circadian rhythms in Drosophila. Cell 99, 791–802 (1999).

17. D. Stoleru, Y. Peng, J. Agosto, M. Rosbash, Coupled oscillators control morning and evening locomotor behaviour of Drosophila. Nature 431, 862–868 (2004).

18. B. Grima, E. Chelot, R. Xia, F. Rouyer, Morning and evening peaks of activity rely on different clock neurons of the Drosophila brain. Nature 431, 869–873 (2004).

19. D. J. Cavanaugh, J. D. Geratowski, J. R. Wooltorton, J. M. Spaethling, C. E. Hector, X. Zheng, E. C. Johnson, J. H. Eberwine, A. Sehgal, Identification of a circadian output circuit for rest:activity rhythms in Drosophila. Cell 157, 689–701 (2014).

20. M. Kunst, M. E. Hughes, D. Raccuglia, M. Felix, M. Li, G. Barnett, J. Duah, M. N. Nitabach, Calcitonin gene-related peptide neurons mediate sleep-specific circadian output in Drosophila. Curr Biol 24, 2652–2664 (2014).

21. F. Guo, J. Yu, H. J. Jung, K. C. Abruzzi, W. Luo, L. C. Griffith, M. Rosbash, Circadian neuron feedback controls the Drosophila sleep--activity profile. Nature 536, 292–297 (2016).

22. S. Yadlapalli, C. Jiang, A. Bahle, P. Reddy, E. Meyhofer, O. T. Shafer, Circadian clock neurons constantly monitor environmental temperature to set sleep timing. Nature 555, 98–102 (2018).

23. F. Guo, M. Holla, M. M. Diaz, M. Rosbash, A Circadian Output Circuit Controls Sleep-Wake Arousal in Drosophila. Neuron 100, 624–635 e624 (2018).

24. L. Sun, R. H. Jiang, W. J. Ye, M. Rosbash, F. Guo, Recurrent circadian circuitry regulates central brain activity to maintain sleep. Neuron 110, 2139–2154 e2135 (2022).

25. H. Dionne, K. L. Hibbard, A. Cavallaro, J. C. Kao, G. M. Rubin, Genetic Reagents for Making Split-GAL4 Lines in Drosophila. Genetics 209, 31–35 (2018).

26. A. Butler, P. Hoffman, P. Smibert, E. Papalexi, R. Satija, Integrating single-cell transcriptomic data across different conditions, technologies, and species. Nat Biotechnol 36, 411–420 (2018).

27. C. Helfrich-Forster, Neurobiology of the fruit fly’s circadian clock. Genes Brain Behav 4, 65–76 (2005).

28. J. D. Ni, A. S. Gurav, W. Liu, T. H. Ogunmowo, H. Hackbart, A. Elsheikh, A. A. Verdegaal, C. Montell, Differential regulation of the Drosophila sleep homeostat by circadian and arousal inputs. eLife 8, (2019).

29. N. Reinhard, E. Bertolini, A. Saito, M. Sekiguchi, T. Yoshii, D. Rieger, C. Helfrich-Forster, The lateral posterior clock neurons of Drosophila melanogaster express three neuropeptides and have multiple connections within the circadian clock network and beyond. The Journal of comparative neurology 530, 1507–1529 (2022).

30. M. H. Alpert, H. Gil, A. Para, M. Gallio, A thermometer circuit for hot temperature adjusts Drosophila behavior to persistent heat. Curr Biol 32, 4079–4087 e4074 (2022).

31. M. Hagemann-Jensen, C. Ziegenhain, P. Chen, D. Ramskold, G. J. Hendriks, A. J. M. Larsson, O. R. Faridani, R. Sandberg, Single-cell RNA counting at allele and isoform resolution using Smart-seq3. Nat Biotechnol 38, 708–714 (2020).

32. X. Long, J. Colonell, A. M. Wong, R. H. Singer, T. Lionnet, Quantitative mRNA imaging throughout the entire Drosophila brain. Nature methods 14, 703–706 (2017).

33. M. E. Hughes, J. B. Hogenesch, K. Kornacker, JTK_CYCLE: an efficient nonparametric algorithm for detecting rhythmic components in genome-scale data sets. Journal of biological rhythms 25, 372–380 (2010).

34. G. Wu, R. C. Anafi, M. E. Hughes, K. Kornacker, J. B. Hogenesch, MetaCycle: an integrated R package to evaluate periodicity in large scale data. Bioinformatics 32, 3351–3353 (2016).

35. B. D. Weger, C. Gobet, F. P. A. David, F. Atger, E. Martin, N. E. Phillips, A. Charpagne, M. Weger, F. Naef, F. Gachon, Systematic analysis of differential rhythmic liver gene expression mediated by the circadian clock and feeding rhythms. Proc Natl Acad Sci U S A 118, (2021).

36. M. Schlichting, S. Richhariya, N. Herndon, D. Ma, J. Xin, W. Lenh, K. Abruzzi, M. Rosbash, Dopamine and GPCR-mediated modulation of DN1 clock neurons gates the circadian timing of sleep. Proc Natl Acad Sci U S A 119, e2206066119 (2022).

37. B. Deng, Q. Li, X. Liu, Y. Cao, B. Li, Y. Qian, R. Xu, R. Mao, E. Zhou, W. Zhang, J. Huang, Y. Rao, Chemoconnectomics: Mapping Chemical Transmission in Drosophila. Neuron 101, 876–893 e874 (2019).

38. M. Kaneko, C. Helfrich-Forster, J. C. Hall, Spatial and temporal expression of the period and timeless genes in the developing nervous system of Drosophila: newly identified pacemaker candidates and novel features of clock gene product cycling. The Journal of neuroscience : the official journal of the Society for Neuroscience 17, 6745–6760 (1997).

39. O. T. Shafer, G. J. Gutierrez, K. Li, A. Mildenhall, D. Spira, J. Marty, A. A. Lazar, M. P. Fernandez, Connectomic analysis of the Drosophila lateral neuron clock cells reveals the synaptic basis of functional pacemaker classes. eLife 11, (2022).

40. Y. Hamasaka, D. Rieger, M. L. Parmentier, Y. Grau, C. Helfrich-Forster, D. R. Nassel, Glutamate and its metabotropic receptor in Drosophila clock neuron circuits. The Journal of comparative neurology 505, 32–45 (2007).

41. M. M. Diaz, M. Schlichting, K. C. Abruzzi, X. Long, M. Rosbash, Allatostatin-C/AstC-R2 Is a Novel Pathway to Modulate the Circadian Activity Pattern in Drosophila. Curr Biol 29, 13–22 e13 (2019).

42. L. S. Mure, H. D. Le, G. Benegiamo, M. W. Chang, L. Rios, N. Jillani, M. Ngotho, T. Kariuki, O. Dkhissi-Benyahya, H. M. Cooper, S. Panda, Diurnal transcriptome atlas of a primate across major neural and peripheral tissues. Science 359, (2018).

43. R. Zhang, N. F. Lahens, H. I. Ballance, M. E. Hughes, J. B. Hogenesch, A circadian gene expression atlas in mammals: implications for biology and medicine. Proc Natl Acad Sci U S A 111, 16219–16224 (2014).

